# Rebound activation of 5-HT neurons following SSRI discontinuation

**DOI:** 10.1101/2023.09.23.558888

**Authors:** Helen M. Collins, L. Sophie Gullino, Dersu Ozdemir, Caroline Lazarenco, Yulia Sudarikova, Elizabeth Daly, Fuencisla Pilar Cuéllar, Raquel Pinacho, David M. Bannerman, Trevor Sharp

**Affiliations:** Dept. of Pharmacology, University of Oxford, UK; Dept. of Experimental Psychology, University of Oxford, UK; Instituto de Biomedicina y Biotecnología de Cantabria, Universidad de Cantabria-CSIC, Spain

## Abstract

Cessation of therapy with a selective serotonin (5-HT) reuptake inhibitor (SSRI) is often associated with an early onset, disabling discontinuation syndrome the mechanism of which is surprisingly little investigated. Here we determined the effect on 5-HT neurochemistry of discontinuation from the SSRI paroxetine. Paroxetine was administered repeatedly to mice (once daily, 12 days versus saline controls) and then either continued or discontinued for up to 5 days. Whereas tissue levels of 5-HT and/or its metabolite 5-HIAA tended to decrease during continuous paroxetine, levels increased above controls after discontinuation, notably in hippocampus. In microdialysis experiments continuous paroxetine elevated hippocampal extracellular 5-HT and this effect fell to saline control levels on discontinuation. However, depolarisation (high potassium)-evoked 5-HT release was reduced by continuous paroxetine but increased above controls post-discontinuation. Extracellular hippocampal 5-HIAA also decreased during continuous paroxetine and increased above controls post-discontinuation. Next, immunohistochemistry experiments found that paroxetine discontinuation increased Fos expression in midbrain 5-HT neurons, adding further evidence for a hyperexcitable 5-HT system. The latter effect was recapitulated by 5-HT_1A_ receptor antagonist administration although gene expression analysis could not confirm altered expression of 5-HT_1A_ autoreceptors following paroxetine discontinuation. Finally, in behavioural experiments paroxetine discontinuation increased anxiety-like behaviour, which partially correlated in time with the measures of increased 5-HT function. In summary, this study finds that SSRI discontinuation triggers a rebound activation of 5-HT neurons across a range of experiments. This effect is reminiscent of neural changes associated with various psychotropic drug withdrawal states, suggesting a common unifying mechanism.

## Introduction

Selective serotonin (5-hydroxytryptamine, 5-HT) reuptake inhibitors (SSRIs) are widely used in the pharmacological treatment of major depression and anxiety-related disorders, as well as a range of other common and disabling neuropsychiatric conditions. Despite their high therapeutic value, abrupt cessation of treatment with an SSRI can produce a debilitating set of psychological and somatic symptoms including heightened anxiety, sleep disruption and sensory disturbances [1, 2]. These symptoms are commonly referred to as a ‘discontinuation’ rather than ‘withdrawal’ syndrome, in part due to SSRIs not being associated with compulsive use, tolerance and craving, and appear within days of treatment cessation. In a recent observational study half of patients experienced the discontinuation syndrome and many reported that their symptoms were severe [3], suggesting a greater clinical problem than previously recognised. This adverse effect of treatment cessation is common to all SSRIs as well as other antidepressants, but the risk of discontinuation syndrome differs between drugs. In particular, paroxetine was estimated to be 10 to 100 times more likely to induce a discontinuation syndrome than other SSRIs [4–6]. Notably, the mechanisms underpinning SSRI discontinuation are currently unknown.

Emerging evidence suggests that SSRI discontinuation occurs across species, which offers opportunities for mechanistic studies. Specifically, rats administered chronic citalopram demonstrated increased startle responsivity within 2 days of discontinuation [7]. In addition, we recently found that mice discontinued from repeated treatment with paroxetine exhibited altered behaviour on the elevated plus maze (EPM) test of anxiety 2 days post-discontinuation [8]. There is a strong evidence linking certain discontinuation symptoms such as anxiety with changes in brain 5-HT [9]. Moreover, since the primary target of SSRIs is the 5-HT transporter, and since 5-HT neurons are well known to adapt to continuous SSRI exposure [10], further changes in 5-HT function during SSRI discontinuation seem likely, as speculated in earlier reviews [11–13]. Surprisingly, however, the effect of SSRI discontinuation on the 5-HT system has been little investigated to date.

In one of the few such studies (that was not in itself directed at discontinuation mechanisms) carried out 30 years ago, continuous administration of fluoxetine exposure caused a decrease in the 5-HT metabolite 5-HIAA in hippocampus and other brain regions [14]. Interestingly, a striking increase in 5-HIAA was evident over 2 weeks following treatment cessation. Similar findings were later observed in mice administered citalopram [15]. It is now well known that continuous SSRI treatment causes an adaptive fall in 5-HT metabolism and synthesis, an effect likely mediated through indirect activation of 5-HT autoreceptors [16–18]. Although changes in 5-HT metabolism/synthesis do not always model 5-HT release [19, 20], these early findings with fluoxetine and citalopram are suggestive of a rebound increase in 5-HT function over days following discontinuation. Notably, evidence of rebound increases in neural function have been observed after withdrawal from various psychotropic drugs including morphine, benzodiazepines and alcohol [21–24], suggesting potential parallels between the mechanisms of cessation of treatment with these different psychotropic drugs.

Against this background the current study investigated the effect of paroxetine discontinuation on 5-HT function in mice using a combination of neurochemical, immunohistochemical and behavioural approaches.

## Methods and Materials

### Animals

C57BL/6J male mice (7 weeks; Charles River) were habituated to the holding facility for one week prior to use. Mice were group housed (3-6 per cage, 21 °C, 12 h light-dark cycle) in cages lined with sawdust bedding and cage enrichment, with *ad libitum* access to food and water. Experiments followed the UK Animals (Scientific Procedures) Act of 1986 and ARRIVE guidelines. We did not use a mixed sex design because in our recent study for reasons female mice did not demonstrate behavioural evidence of SSRI discontinuation [8]. This possibly reflects recent clinical findings that compared to women men were more likely to experience discontinuation symptoms, and symptoms are more likely to be severe [5].

### Drug treatment

Mice were allocated to treatment groups by stratified randomisation. Mice received once-daily injections of either 10 mg/kg s.c. paroxetine (1mg/ml; Abcam) or saline vehicle for 12 days, then treatment was either continued or swapped to saline (discontinuation groups) for a further 2 or 5 days [8]. In some experiments (*ex vivo* neurochemistry, immunohistochemistry), 90 min prior to brain removal mice were tested on the elevated plus maze (EPM) to provide behavioural measures to correlate with the neurochemical effects of SSRI discontinuation.

### *Ex vivo* 5-HT neurochemistry

After cervical dislocation brains were rapidly removed and frozen in isopentane on dry ice, prior to storage at −80 °C. Subsequently, hippocampus, striatum, frontal cortex and midbrain were dissected on ice, weighed and then placed in 0.09 M perchloric acid prior to sonication and centrifugation (13,000 rpm for 15 min, 4 °C). Supernatant 5-HT and 5-HIAA, as well as dopamine and its metabolite DOPAC, were measured using high performance liquid chromatography (HPLC) with electrochemical detection [25].

### *In vivo* microdialysis

Mice were anaesthetised with urethane (1 g/kg *i.p.*) and maintained at 36 ± 1 °C using a homeothermic blanket. A microdialysis probe (2 mm, Microbiotech MAB4) was stereotaxically implanted into the hippocampus (AP −3.0 mm, ML ±2.9 mm, DV −4.0 mm relative to bregma and the dura surface; [26]) and perfused (2 µl/min) with artificial cerebrospinal fluid (in mM: 140 NaCl, 4 KCl, 1.2 Na_2_HPO_4_, 0.27 NaH_2_PO_4_, 1 MgCl_2_, 2.4 CaCl_2_ and 7.2 glucose). After a 60 min post-implantation period, samples were collected every 20 min for 120 min. The perfusion medium was then switched (20 min) to one containing 56 mM KCl, followed 60 min later by 100 mM KCl. Microdialysate samples were analysed for 5-HT and 5-HIAA using HPLC with electrochemical detection (see above).

### Fos immunohistochemistry

Ninety min after EPM exposure (see below) mice were injected with pentobarbital (200 mg/kg *i.p.*) and transcardially perfused with phosphate buffered saline (PBS) followed by 4 % paraformaldehyde (PFA) in PBS. Brains were kept at 4 °C in 4 % PFA for 48 h, then stored in cryoprotective 30 % sucrose in PBS at 4 °C. Cryostat cut sections (30 μm, coronal) containing the dorsal raphe nucleus (DRN) were sequentially washed in PBS, ammonium chloride and PBS with 0.3 % TWEEN® 80 (PBS-T). Sections were then incubated in PBS-T with 10 % donkey serum for 1 h at room temperature, and then incubated overnight at 4 °C with the primary antibodies for c-Fos (Abcam, ab214672), TPH2 (Abcam, Ab121013) and NeuN (Abcam, Ab104224) in PBS-T with 2 % donkey serum. Sections were then washed in PBS-T and incubated for 2 h (room temperature) with donkey anti-rabbit IgG (Alexa Flor™ 488; Invitrogen, A21206), donkey anti-goat IgG (Alexa Flor™ 568; Invitrogen, A11057) and donkey anti-mouse IgG (Alexa Flor™ 647; Invitrogen, A21202) in PBS-T with 2 % donkey serum. After final washes in PBS-T then PBS, sections were mounted on glass slides and imaged (Olympus Epifluorescence Microscope BX40 with ImageJ Micromanager v1.4). c-Fos/TPH2 double-labelled DRN neurons were counted in three sections per mouse, averaged, and expressed as the number of neurons per mm^2^. All counting was conducted blind to treatment.

### qPCR analysis

For PCR analysis, midbrain raphe region and frontal cortex were rapidly dissected from frozen tissue sections (1 mm). RNA extraction, cDNA conversion and qPCR were conducted as described [27]. In brief, RNA was extracted (Qiagen RNeasy Mini Kit) and eluted into RNase-free water (20 μl for midbrain, 25 μl for cortex) prior to conversion to cDNA using a high-capacity cDNA Reverse Transcription Kit (Life Technologies) and T100 Thermocycler (Bio-Rad). QPCR was performed (200 ng RNA) using a LightCycler® 480 instrument (Roche Diagnostics) with the following primers (5’ to 3’ at 300 nM): *5-HT_1A_* forward GACAGGCGGCAACGATACT, reverse CCAAGGAGCCGATGAGATAGTT [28]; *5-HT_1B_* forward CCCATCAGCACCATGTACAC, reverse GACTTGGTTCACGTACACAG [29]; *5-HT_2A_* forward CAGGCAAGTCACAGGA TAGC, reverse TTAAGCAGAAAGAAAATCCCACAG [30]; *5-HT_4_* forward CCTCACAGCAACTTCTCCTT, reverse TCCCCTGACTTCCTCAAATA [31]. β-actin was the housekeeping gene; forward CATTGCTGACAGGATGCAGAAGG, reverse TGCTGGAAGGTGGACAGTGAGG [32]. Reactions (384 well-plates, 10 μl reaction volume, 5 μl Precision®PLUS qPCR Master Mix with SYBRgreen, 25 ng cDNA) used the following cycle: enzyme activation for 2 min at 95 °C, 40 cycles of 10 s at 95 °C, 1 min at 60 °C, then held at 4 °C. Data were analysed using ΔCt values with outliers identified by ROUT analysis of 2^-ΔΔCt^ values.

### Elevated plus maze

EPM experiments were performed as previously described [8]. In brief, experiments were conducted in the light phase (10:00–15:00 h) by an observer blind to treatment. The EPM (50 cm above the floor) comprised 2 open arms (35×6 cm) perpendicular to 2 closed arms (35×6 cm, 20 cm walls) with a central region (6×6 cm), placed in a dimly lit room. Mice were placed facing the walls of the closed arm (counterbalanced between the 2 closed arms) and movement was automatically tracked for 300 s (ANY-maze software, Stoelting Co.). Head dips were counted manually.

### Statistical analysis

Data were initially assessed for normality with D’Agostino Pearson’s test. Multi-time point microdialysis data were analysed using repeated measures ANOVA with Bonferroni’s multiple comparisons test. Other parametric data were analysed by one-way ANOVA with post-hoc Fisher’s Least Significant Difference (LSD). The Kruskal Wallis test with Uncorrected Dunn’s test was used for non-parametric data. ROUT analysis was used to identify outliers in all datasets. Data were analysed with GraphPad Prism (v9) by an experimenter blind to treatment group. P<0.05 was considered statistically significant.

## Results

### Paroxetine discontinuation increased 5-HT metabolism in hippocampus ex vivo

Initial experiments investigated whether discontinuation from paroxetine caused a rebound increase in 5-HT metabolism in hippocampus and other brain regions as previously reported for fluoxetine [14]. For hippocampus, continued paroxetine tended to reduce levels of 5-HIAA and 5-HT compared to saline controls, but this effect recovered on discontinuation day 2 and increased above control levels on discontinuation day 5 (5-HIAA, F_(3,47)_=4.626, p=0.0065; 5-HT, F_(3,47)_=5.632, p=0.0022; for post hoc analysis see Fig. 1A). Other regions showed similar trends to hippocampus although statistically the most consistent finding was 5-HT and 5-HIAA levels were higher on discontinuation day 5 compared to continued paroxetine; striatum (5-HIAA, F_(3,47)_=3.529, p=0.0218; 5-HT, F_(3,47)_=2.820, p=0.0490; Fig. 1B), frontal cortex (5-HIAA, F_(3,47)_=4.435, p=0.0082; 5-HT, F_(3,47)_=2.722, p=0.0551; Fig. 1C) and midbrain (5-HIAA, F_(3,47)_=5.305, p=0.0031; 5-HT, F_(3,47)_=4.609, p=0.0066; Fig. 1D).

**Figure 1.**
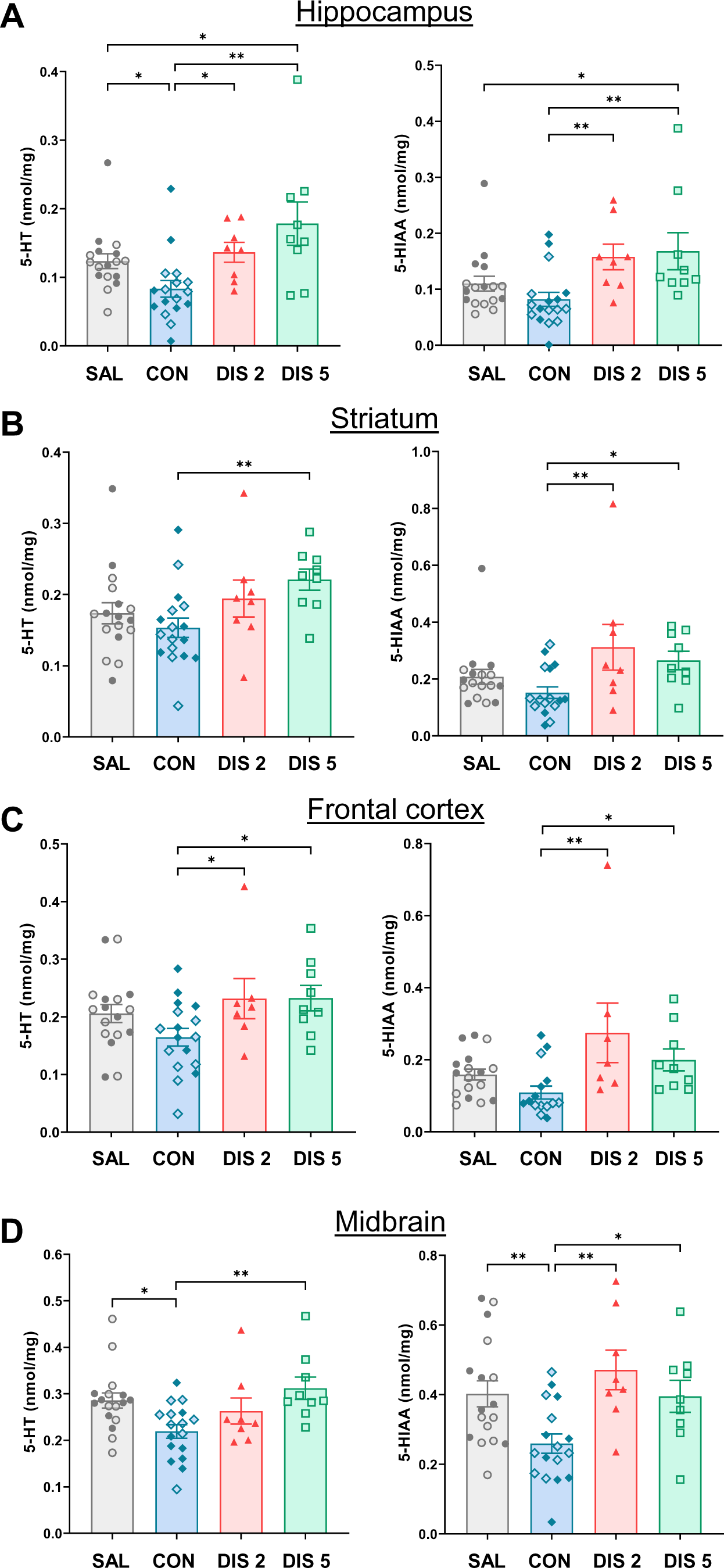
Effect paroxetine discontinuation on tissue levels of 5-HT and 5-HIAA in (A) hippocampus, (B) striatum, (C) frontal cortex and (D) midbrain. Mice received either saline (SAL) or 10 mg/kg *s.c.* paroxetine (CON) once daily for 12 days and then paroxetine was discontinued for 2 days (DIS 2) or 5 days (DIS 5). Each column is a mean ± SEM value of the individual points shown. Values for saline and continuous paroxetine groups were pooled across experiments (open and closed symbols are matched across experiments). Data were analysed using one-way ANOVA with post hoc Fisher’s LSD, *p<0.05, **p>0.01.

Compared to saline controls, neither continuous treatment with paroxetine nor 2 or 5 days of discontinuation altered tissue levels of DOPAC or dopamine in any brain region (Tables S1 and S2).

### Paroxetine discontinuation decreased extracellular 5-HT but increased 5-HT metabolism in hippocampus in vivo

Next, we investigated the effect of paroxetine discontinuation on basal extracellular 5-HT and 5-HIAA in hippocampus using *in vivo* microdialysis (Fig. 2A). Mice receiving continuous paroxetine had 2-3 times higher levels of extracellular 5-HT than saline controls (Fig. 2B). This effect rapidly reversed on discontinuation from paroxetine, and basal extracellular 5-HT fell to saline controls at discontinuation days 2 and 5 (F_(3,28)_=9.229, p=0.0002; for post hoc analysis see Fig. 2B).

**Figure 2.**
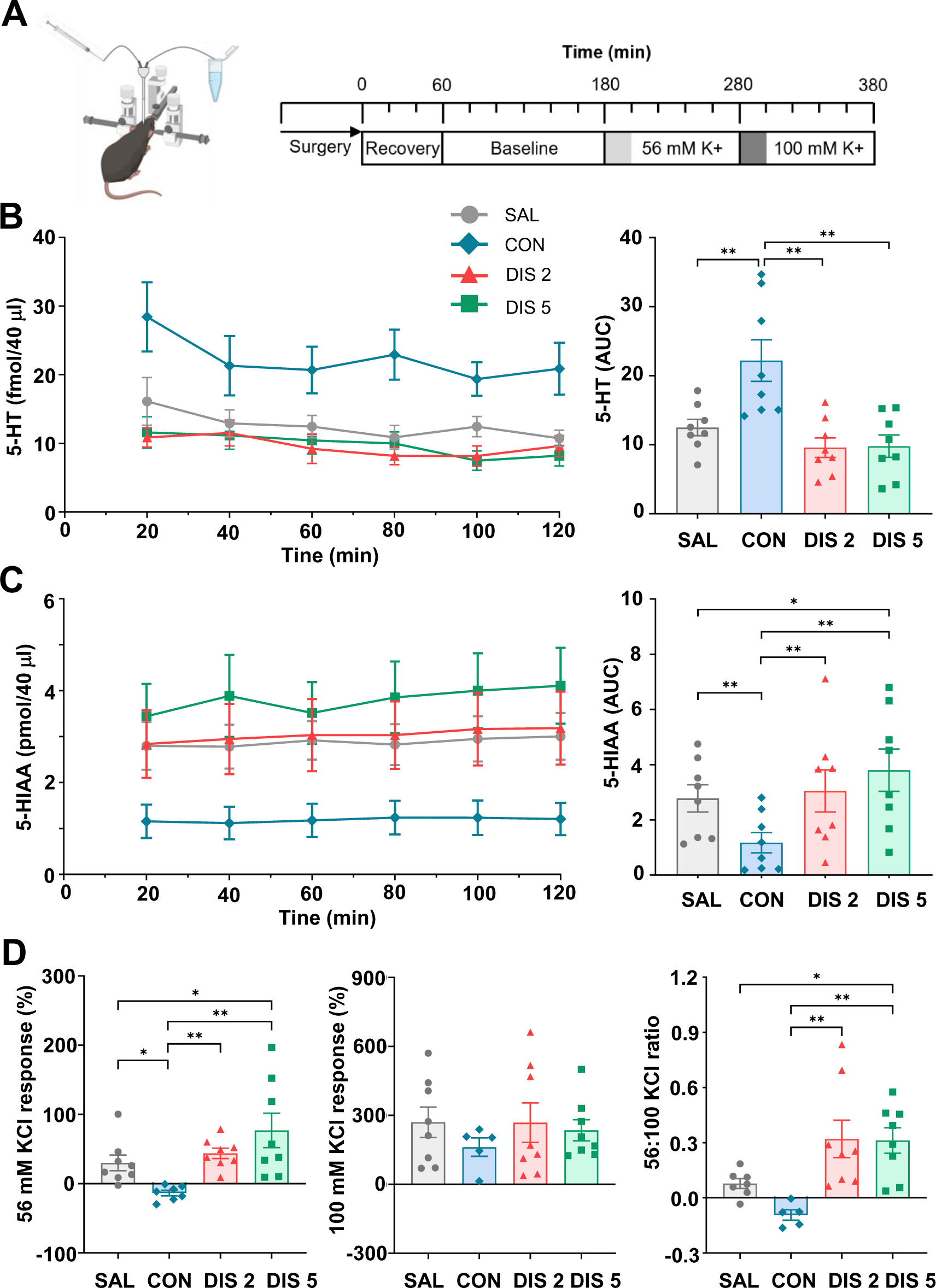
Effect of paroxetine discontinuation on extracellular 5-HT and 5-HIAA in hippocampus as measured by microdialysis in anaesthetised mice. Mice received either saline (SAL) or 10 mg/kg *s.c.* paroxetine (CON) once daily for 12 days and then paroxetine was discontinued for 2 days (DIS 2) or 5 days (DIS 5). Mice were then subject to the experimental design illustrated (A). Baseline levels of 5-HT (B) and 5-HIAA (C) at individual time points (left) and averaged over the time course (right). (D) Effect of perfusion with 56 mM KCl (left) or 100 mM KCl (middle) together with 100 mM: 56 mM KCl ratio (right). Mean ± SEM values are shown. Data analysed using either repeated measures ANOVA with post-hoc Bonferroni’s multiple comparisons test (time course) or one-way ANOVA with Fisher’s LSD (time course averages), * p<0.05, ** p<0.01.

In comparison, basal extracellular levels of 5-HIAA were 40-50 % lower in mice receiving continuous paroxetine compared to saline controls. This effect had reversed on day 2 after discontinuation and on day 5 levels of 5-HIAA were ∼30 % above saline controls (F_(3,28)_=10.165, p<0.0001; for post hoc analysis see Fig. 2C).

Thus, continuous paroxetine increased basal extracellular 5-HT in hippocampus and this effect fell rapidly to saline control levels following discontinuation. However, on discontinuation hippocampal extracellular 5-HIAA showed a rebound increase as detected with *ex vivo* measurements.

### Paroxetine discontinuation increased depolarisation-evoked 5-HT release in hippocampus in vivo

Basal extracellular levels of 5-HT reflect a combination of processes including 5-HT reuptake, diffusion, synthesis, metabolism and release. Local perfusion with high KCl was used to measure depolarisation-evoked 5-HT release. In saline controls 56 mM KCl caused a short-lasting increase in 5-HT of ∼30 %. This response was attenuated in mice receiving continuous paroxetine, which reversed on discontinuation day 2, and was greater than saline controls on discontinuation day 5 (F_(3,27)_=6.198, p=0.0024; for post hoc analysis see Fig. 2D).

In comparison, 100 mM KCl evoked an increase in 5-HT of ∼ 250 % in saline controls, and this effect was not different across the treatment groups (F_(3,25)_=0.4583, p=0.7139; Fig. 2D). Using the 56:100 mM KCl response ratio as an overall measure of sensitivity to high KCl, a rebound increase in depolarisation-evoked 5-HT was evidence on both discontinuation days 2 and 5 (F_(3,24)_=6.642, p=0.002; for post hoc analysis see Fig. 2D). Collectively, these findings suggest that 5-HT terminals in the hippocampus were more sensitive to depolarisation on days 2 and 5 following paroxetine discontinuation.

### Paroxetine discontinuation increased c-Fos expression in 5-HT neurons

The above findings suggest that 5-HT neurons are more excitable during paroxetine discontinuation. To investigate this further, co-localisation of Fos and TPH2 was used as a marker of 5-HT neuron activity. Multiple c-Fos/TPH2 co-labelled neurons were observed in the DRN (Fig. 3A). Interestingly, on both discontinuation day 2 and 5 the number of c-Fos/TPH2 co-labelled neurons was increased compared to continuous paroxetine and saline controls (one-way ANOVA: day 2, F_(2,16)_=5.637, p=0.0140; day 5, F_(2,15)_=5.523, p=0.0159; for post hoc analysis see Fig. 3B). Compared to saline controls the number of c-Fos/TPH2 co-labelled DRN neurons had a tendency to be reduced by continuous paroxetine although this effect was not statistically significant (Fig. 3B).

**Figure 3.**
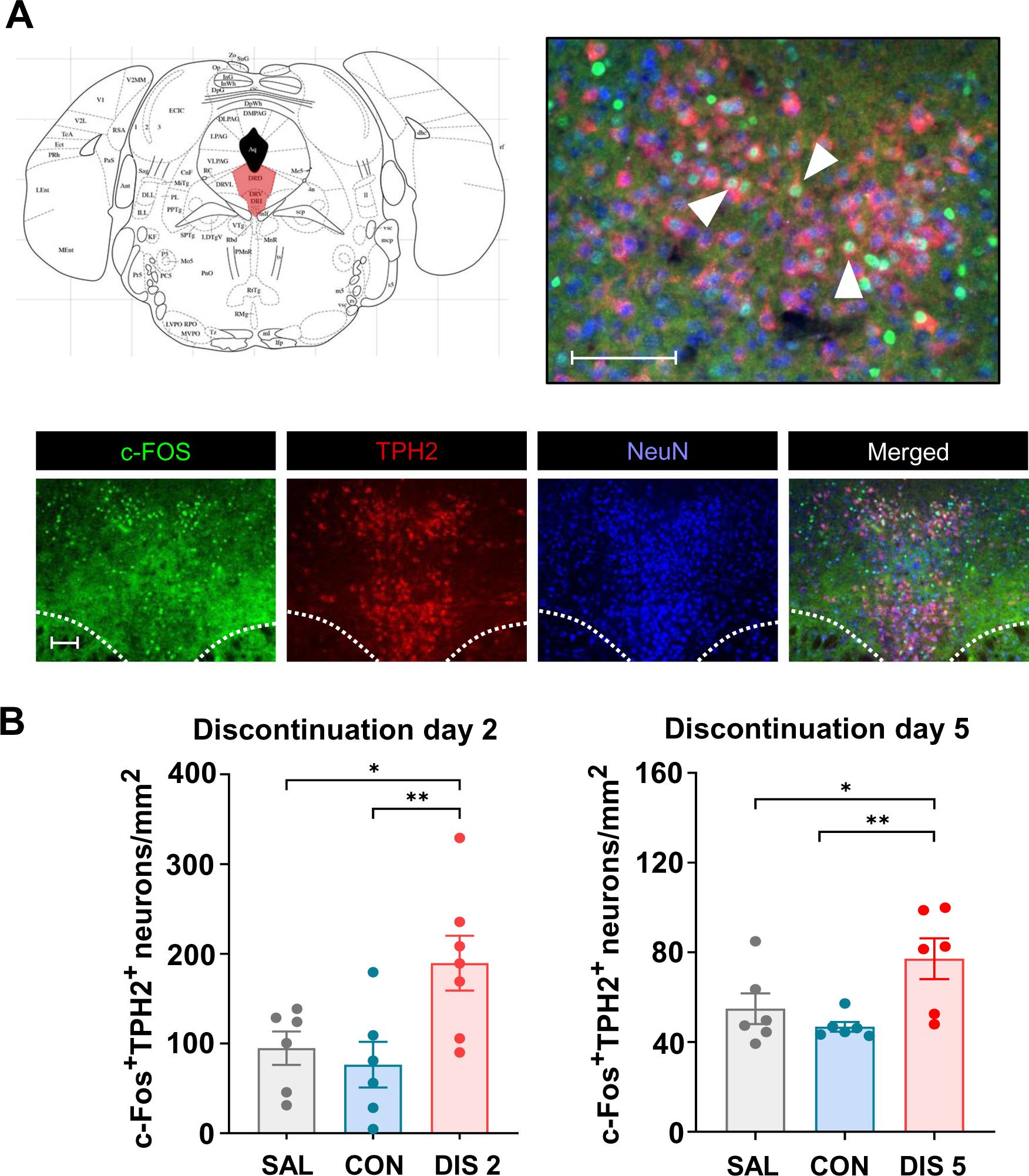
Effect of paroxetine discontinuation on Fos/TPH2 double-labelled neurons in the DRN. Mice received either saline (SAL) or 10 mg/kg *s.c.* paroxetine (CON) once daily for 12 days and then paroxetine was discontinued for 2 days (DIS 2) or 5 days (DIS 5). (A) Illustration of DRN localisation (red) together with images of c-Fos, TPH2 and NeuN immunoreactivity and their merger at low (x10) and high (x40) resolution (scale bar 100 μm; white arrows show Fos/TPH2 co-labelled neurons). Dotted lines indicate the medial longitudinal fasciculi. (B) Quantified data showing number of showing Fos/TPH2 co-labelled neurons. Mean ± SEM values are shown. Data analysed using one-way ANOVA with post-hoc Fisher’s LSD, *p<0.05, ** p<0.01.

This finding of increased c-Fos/TPH2 co-labelled neurons in the DRN of mice discontinued from paroxetine further supports the idea that 5-HT neurons were more excitable.

### Possible role of 5-HT feedback in effects of paroxetine discontinuation

A possible mechanism underlying hyperexcitable 5-HT neurons is the state of 5-HT_1A_ autoreceptor desensitization which may have been exposed on cessation of paroxetine treatment. PCR analysis investigated whether paroxetine discontinuation was associated with adaptive changes in 5-HT_1A_ autoreceptor expression, as well as the expression of other 5-HT receptor subtypes in the midbrain raphe region and prefrontal cortex linked to the feedback control of DRN 5-HT neurons (e.g. [33]).

On discontinuation day 2 mice had reduced 5-HT_1A_ mRNA expression in both the midbrain raphe region and frontal cortex compared to continued paroxetine treatment (see Table S3 for statistical details) but expression of the 5-HT_1B_ receptor in the midbrain raphe region or 5-HT_2A_ and 5-HT_4_ receptors in frontal cortex were unchanged (Table S3). On discontinuation day 5, however, there was no effect of treatment on the expression of any 5-HT receptor subtype in either the midbrain raphe region or frontal cortex (Table S3).

To further test whether changes in 5-HT feedback may be involved in discontinuation responses, we tested the effect of the selective 5-HT_1A_ receptor antagonist WAY-100635 (1 mg/kg s.c.) on c-Fos expression. WAY-100635 increased the number of c-Fos/TPH2 co-labelled neurons compared to saline-controls (167.3±6.0 versus 95.7±13.6 neurons/mm^2^; t_(10)_=3.118, p=0.0109). This result adds to the plausibility that loss of 5-HT_1A_ receptor feedback control contributed to increased excitability of 5-HT neurons on paroxetine discontinuation.

### Paroxetine discontinuation increased anxiety-like behaviour on the EPM

Finally, given the well-established link between increased 5-HT and anxiety-like behaviour [9, 34], experiments tested the temporal relationship between the discontinuation-evoked increase in 5-HT function and changes in anxiety-like behaviour as assessed by performance on the EPM.

On discontinuation day 2 mice spent less time in the open arms (F_(2,33)_=9.902, p=0.0004), made fewer open arm entries (F_(2,33)_=7.708, p=0.0018) and showed fewer head dips (F_(2,33)_=11.29, p=0.0002) compared to mice receiving continued paroxetine or saline (for post hoc analysis see Fig. 4B). These mice also had reduced distance travelled on the EPM (F_(2,33)_=7.415, p=0.0022; Fig. 4B), supporting the notion of increased behavioural inhibition in the anxiogenic environment. In contrast, on discontinuation day 5 the performance of mice on the EPM was not different from continued paroxetine or saline controls (open arm time: F_(2,33)_=1.451, p=0.2490; open arm entries: F_(2,33)_=1.100, p=0.3449; head dips: F_(2,33)_=2.422, p=0.1043; distance: F_(2,33)_=2.588, p=0.0908; for post hoc analysis see Fig. 4C).

**Figure 4.**
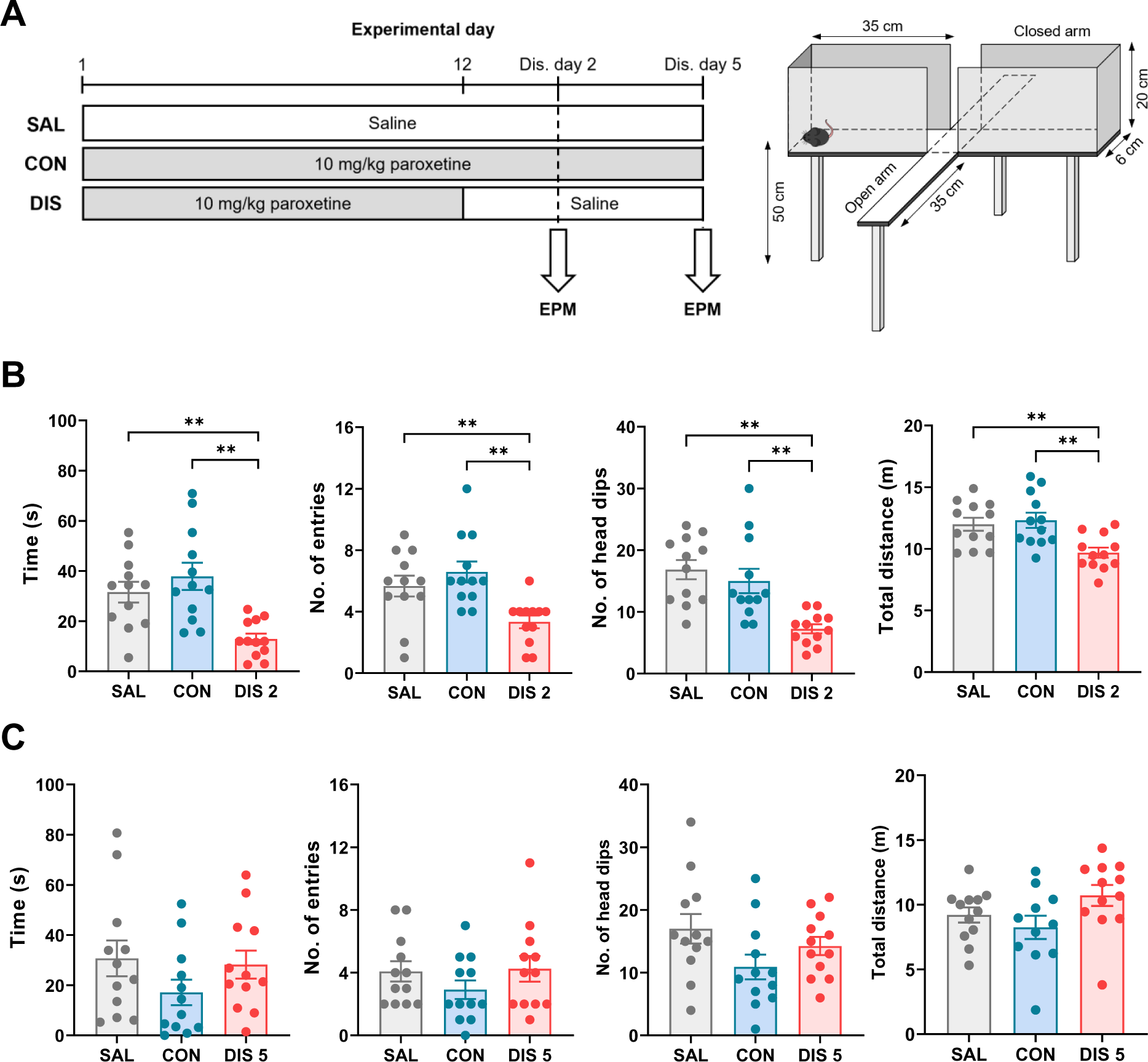
Effect of paroxetine discontinuation on anxiety-like behaviour on the EPM. Mice received either saline (SAL) or 10 mg/kg *s.c.* paroxetine (CON) once daily for 12 days and then paroxetine was discontinued for 2 days (DIS 2) or 5 days (DIS 5). (A) Experimental design with illustration of EPM apparatus. (B) EPM parameters for day 2 of discontinuation. (C) EPM parameters for day 5 of discontinuation. Mean ± SEM values are shown. Data analysed using one-way ANOVA with Fisher’s LSD, ** p<0.01.

Overall, these data indicate that the rebound increase in 5-HT function observed on discontinuation day 2 was associated with increased anxiety-like behaviour. However, this temporal correlation was not apparent on discontinuation day 5.

## Discussion

Abrupt cessation of a course of treatment with an SSRI is often associated with a disabling discontinuation syndrome (see Introduction), which is currently of unknown cause and little studied. Here we found that across experiments, abrupt cessation of treatment with paroxetine was associated with evidence of increased 5-HT neuronal function. Specifically, paroxetine discontinuation; i) increased tissue levels of 5-HIAA and 5-HT, especially in hippocampus *ex vivo* (without altering dopamine or its metabolite DOPAC), ii) increased extracellular 5-HIAA in hippocampus *in vivo*, iii) increased depolarisation-evoked release of hippocampal 5-HT *in vivo*, and iv) increased c-Fos immunoreactivity in 5-HT neurons (TPH2-immunolabelled) in the DRN *ex vivo*. These effects occurred during a 5 day discontinuation period, with some (evoked 5-HT release, increased c-Fos expression) being already detected after 2 days. Finally, several experiments observed a predicted decrease in 5-HT neuronal function in mice receiving continued paroxetine that preceded the rebound increase.

The emergence of these discontinuation effects is consistent with the short half-life of paroxetine in rodents (t_1/2_=6.3 h, [35]) as well as paroxetine having non-linear metabolism kinetics and inhibiting its own metabolising enzyme (CYP2D6) such that when plasma paroxetine levels fall its metabolism increases and its washout is accelerated [36]. The timing also fits with behavioural effects appearing within 2 days of paroxetine discontinuation as observed in the present and a recent study in animals [8], as well as previous studies in depressed patients [37, 38]. Moreover, earlier studies reported increased 5-HT metabolism/synthesis in rodent hippocampus and other brain regions within days of discontinuation from fluoxetine or escitalopram [14, 15]. Here, the rebound increase in 5-HT neuron function was not investigated beyond 5 days after discontinuation but the latter studies on 5-HT metabolism/synthesis suggest that the rebound effect may last several weeks.

A plausible explanation for the rebound increase in 5-HT transmission following paroxetine cessation is removal of the constant suppression of 5-HT neurons produced during repeated exposure to the drug, mediated by negative 5-HT feedback mechanisms. That these feedback mechanisms were operational during continuous paroxetine was evident both as reduced 5-HT metabolism and reduced depolarisation-evoked release of 5-HT in hippocampus. Previous studies also report reduced brain 5-HT synthesis and metabolism in rodents chronically exposed to paroxetine and other SSRIs [17, 18]. The negative feedback effects of continuous SSRI administration likely result from the inhibition of 5-HT uptake causing a sustained elevation in extracellular 5-HT. Here, continuous paroxetine increased extracellular 5-HT 2-3 fold in hippocampus in accord with previous findings in this and other brain regions including the DRN [39]. Increased extracellular 5-HT will activate various 5-HT feedback mechanisms including somatodendritic 5-HT_1A_ autoreceptors and terminal 5-HT_1B_ autoreceptors, and potentially 5-HT receptors located on postsynaptic neurons that suppress the firing of 5-HT neurons [33]. Indeed, SSRI-induced decreases in 5-HT neuronal activity and synthesis/metabolism can be attenuated by 5-HT_1A/1B_ receptor antagonists [18, 40]. Hence, a likely scenario is that on abrupt cessation of SSRI administration and reversal of 5-HT transporter inhibition, extracellular 5-HT rapidly falls (Fig. 2B) relieving inhibitory 5-HT feedback systems, which triggers a rebound increase in 5-HT synthesis, release and neuronal excitability.

The present finding of reduced 5-HT_1A_ receptor mRNA in the midbrain raphe region and frontal cortex at 2 days after paroxetine discontinuation comprises evidence that loss of 5-HT feedback control contributes to the rebound increase in 5-HT transmission. This change was not evident at 5 days but 5-HT_1A_ autoreceptor desensitization during continuous SSRI administration has been difficult to demonstrate at the gene/protein expression, as opposed to functional, level [41–43]. Previously, mice with a genetic depletion of somatodendritic 5-HT_1A_ autoreceptors demonstrated increased firing of 5-HT neurons and increased physiological reactivity to stress [44]. Moreover, here administration of the selective 5-HT_1A_ receptor antagonist WAY-100635 mimicked the effect of paroxetine discontinuation on c-Fos/TPH2 double-labelled DRN neurons, in accord with previous studies showing that WAY-100635 increased the firing of 5-HT neurons [45, 46]. This is not to say that 5-HT_1A_ receptor blockade fully models the effects of SSRI discontinuation, not least because such agents block not only pre-but also postsynaptic 5-HT_1A_ receptors. Nevertheless, rapid relief from negative 5-HT feedback control seems a likely contributor to an increase in excitability of 5-HT neurons after SSRI discontinuation although the present study does not rule out the involvement of other adaptive changes.

A rebound increase in 5-HT seems difficult to reconcile with the fact that reinstatement of SSRI treatment is often used to manage the discontinuation. However, re-introduction of an SSRI would likely increase extracellular 5-HT and reduce 5-HT neuron firing by indirectly activating inhibitory feedback mechanisms such as somatodendritic 5-HT_1A_ autoreceptors. In effect, recommencement of SSRI treatment would restore 5-HT neuron feedback and thereby dampen down 5-HT neuron hyperexcitability that is a potential driver of discontinuation symptoms. The idea of increased in excitability of 5-HT neurons also contrasts with earlier speculation that SSRI discontinuation may be mediated by *reduced* synaptic 5-HT due to rapid relief of 5-HT transporter blockade [11]. The latter idea was largely based on the observation that the symptomatic effects of tryptophan depletion in SSRI-treated patients resembled those of SSRI discontinuation. However, others have suggested that the effects of tryptophan depletion were more characteristic of depression relapse than those of SSRI discontinuation [47]. On the other hand, it is conceivable that during continuous SSRI exposure, an acute fall in 5-HT induced by tryptophan depletion relieves 5-HT feedback which may generate 5-HT neuron instability; in this case effects of tryptophan depletion in SSRI-treated patients may indeed reflect SSRI discontinuation.

The current study found that day 2 following paroxetine discontinuation, mice showed increased anxiety-like behaviour on the EPM. This replicates the finding in our recent study [8] and is consistent with an earlier report of increased acoustic startle responsivity in rats within a few days of discontinuation from citalopram [7]. There is much evidence that increased 5-HT transmission generates an anxiogenic effect on the EPM and in other anxiety paradigms [9, 34], suggesting that the discontinuation-induced increase in 5-HT function and anxiety-like behaviour may be causally linked. However, the anxiogenic effect of paroxetine discontinuation had dissipated by discontinuation day 5 when increases in 5-HT neuronal function were still evident. Explanations for this mismatch in timing include the possibility that there is adaptation to increased 5-HT function on discontinuation day 2, resulting in normalisation of behaviour on the EPM. It is also possible that other mechanisms contribute to increased anxiety-like behaviour. For example, it is speculated that a rebound increase in cholinergic transmission contributes to discontinuation effects of various antidepressants including paroxetine, which has moderate affinity for muscarinic receptors [11, 12, 48].

Aside from anxiety, a rebound increase in 5-HT neurotransmission after paroxetine discontinuation might generate other behaviours or physiological changes that were not monitored here. For instance, we recently reported that mice demonstrated evidence of sleep disruption that commenced 2 days after discontinuation from paroxetine and continued for up to 9 days [49]. Sleep is well known to be regulated by 5-HT, and sleep disturbances are a recognised feature of SSRI discontinuation syndrome.

Interestingly, increased 5-HT neuron excitability in response to SSRI discontinuation has parallels with a previously proposed account of withdrawal from psychotropic drug administration. According to this account, adaptive influences develop when neural systems are subjected to prolonged suppression and rebound when, on drug removal, these oppositional influences no longer meet resistance [23]. Thus, rebound increases in neurotransmission are reported in response to discontinued administration of other psychotropic drugs; for example, a rebound increase in excitatory transmission is associated with alcohol and benzodiazepine withdrawal [21, 24] whereas opiate withdrawal is associated with elevated noradrenergic activity [22]. Variation in the symptoms of withdrawal from these different classes of psychotropic drugs likely reflects the different neurotransmitter systems involved [23].

In conclusion, we report evidence that SSRI discontinuation is associated with a rebound increase in 5-HT function that lasts many days. We speculate that this effect is a consequence of a rapid fall in extracellular 5-HT following abrupt cessation of SSRI administration, leading to disinhibition of 5-HT neurons via removal of inhibitory 5-HT feedback. This response to SSRI discontinuation resembles the changes of other neural systems following cessation of treatment with other psychotropic drugs, suggesting a common neurobiological mechanism.

## Author contributions

HMC performed experiments, analysed the data, and contributed to writing the manuscript. FPC and SG performed the qPCR experiments. YS, ED and DO contributed to the *ex vivo* tissue analysis. CL contributed to the EPM experiments. RP aided design of the work and manuscript preparation. TS and DMB made contributions to the conception and design of the work, drafting and revising the manuscript and analysis and interpretation of data.

