## Supplementary for "Rebound activation of 5-HT neurons following SSRI discontinuation"

**Table S1 Effect of 2 days discontinuation from 12 days once-daily paroxetine (10 mg/kg) on regional brain tissue levels of DA and DOPAC.** Mean± SEM values are shown (nmol/mg tissue), 8 mice/group. Repeated measures ANOVA with Bonferroni's test. DA: effect of treatment $F(2,21)=0.844$, $p=0.433$, effect of region $F(3,63)=74.49$, $p<0.001$, treatment*region interaction $F(6,63)=0.263$, $p=0.953$. DOPAC: effect of treatment $F(2,21)=0.364$, $p=0.696$, effect of region $F(3,63)=51.21$, $p<0.001$, treatment*region interaction $F(6,63)=0.256$, $p=0.956$.

| | Treatment | | |
|---|---|---|---|
| **Brain region** | **SAL** | **CON** | **DIS2** |
| DA | | | |
| Midbrain | 0.111 ± 0.016 | 0.135 ± 0.014 | 0.134 ± 0.025 |
| Hippocampus | 0.132 ± 0.021 | 0.162 ± 0.032 | 0.148 ± 0.029 |
| Striatum | 3.025 ± 0.529 | 3.448 ± 0.415 | 3.514 ± 0.610 |
| Frontal cortex | 0.657 ± 0.114 | 0.921 ± 0.140 | 1.219 ± 0.493 |
| DOPAC | | | |
| Midbrain | 0.030 ± 0.002 | 0.026 ± 0.003 | 0.031 ± 0.006 |
| Hippocampus | 0.020 ± 0.004 | 0.020 ± 0.007 | 0.021 ± 0.002 |
| Striatum | 0.289 ± 0.067 | 0.330 ± 0.053 | 0.309 ± 0.051 |
| Frontal cortex | 0.120 ± 0.022 | 0.135 ± 0.011 | 0.174 ± 0.047 |

**Table S2 Effect of 5 days discontinuation from 12 days once-daily paroxetine (10 mg/kg) on regional brain tissue levels of DA and DOPAC.** Mean± SEM values are shown (nmol/mg tissue) with 9 mice/group. Repeated measures ANOVA with Bonferroni's test. N=9/group. DA: effect of treatment $F(2,24)=2.325$, $p=0.109$, effect of region $F(3,72)=15.00$, $p<0.001$, treatment*region interaction $F(6,72)=0.539$, $p=0.708$. DOPAC: effect of treatment $F(2,24)=2.483$, $p=0.092$, effect of region $F(6,72)=71.07$, $p<0.001$, treatment*region interaction $F(6,72)=0.827$, $p=0.554$.

| **Brain region** | **Treatment**<br>**SAL** | <br>**CON** | <br>**DIS5** |
|---|---|---|---|
| DA | | | |
| Midbrain | 0.048 ± 0.010 | 0.063 ± 0.15 | 0.067 ± 0.013 |
| Hippocampus | 0.074 ± 0.025 | 0.084 ± 0.025 | 0.163 ± 0.047 |
| Striatum | 0.975 ± 0.271 | 1.935 ± 0.399 | 2.190 ± 0.498 |
| Frontal cortex | 0.419 ± 0.115 | 0.423 ± 0.142 | 0.613 ± 0.128 |
| DOPAC | | | |
| Midbrain | 0.025 ± 0.004 | 0.028 ± 0.004 | 0.056 ± 0.034 |
| Hippocampus | 0.013 ± 0.004 | 0.020 ± 0.003 | 0.033 ± 0.008 |
| Striatum | 0.291 ± 0.063 | 0.342 ± 0.058 | 0.391 ± 0.044 |
| Frontal cortex | 0.073 ± 0.017 | 0.086 ± 0.023 | 0.128 ± 0.025 |

**Table S3 Effect of 2 and 5 days of paroxetine discontinuation on mRNA expression of 5-HT receptor subtypes in mouse midbrain raphe region and prefrontal cortex.** Mice were injected once daily for 12 days with either saline (SAL) or 10 mg/kg paroxetine (CON), with the latter discontinued for either 2 (DIS 2) or 5 (DIS 5) days. Values for mRNA (mean ± SEM) are relative to house-keeping genes, n=6-8/group. One-way ANOVA - post-hoc comparisons, # p<0.05 CON vs DIS, * p<0.01 CON vs DIS.

| | Discontinuation day 2 | | | |
|---|---|---|---|---|
| | **SAL** | **CON** | **DIS 2** | **ANOVA** |
| Midbrain region | | | | |
| *5-HT1A* | 1.03 ± 0.10 | 1.34 ± 0.13 | 0.99 ± 0.10# | F(2,21)=2.949, p=0.0444 |
| *5-HT1B* | 1.05 ± 0.13 | 1.23 ± 0.20 | 0.88 ± 0.14 | F(2,21)=1.417, p=0.2835 |
| PFC | | | | |
| *5-HT1A* | 1.02 ± 0.08 | 1.10 ± 0.10 | 0.76 ± 0.07* # | F(2,21)=4.529, p=0.0232 |
| *5-HT1B* | 1.05 ± 0.11 | 1.88 ± 0.42 | 1.44 ± 0.23 | F(2,21)=2.098, p=0.1477 |
| *5-HT2A* | 1.01 ± 0.06 | 1.03 ± 0.09 | 0.93 ± 0.10 | F(2,21)=0.3668 p=0.6973 |
| *5-HT4* | 1.08 ± 0.16 | 0.94 ± 0.09 | 1.81 ± 0.64 | F(2,21)=1.264 p=0.3052 |

| | Discontinuation day 5 | | | |
|---|---|---|---|---|
| | **SAL** | **CON** | **DIS 5** | **ANOVA** |
| Midbrain region | | | | |
| *5-HT1A* | 1.03 ± 0.12 | 0.85 ± 0.06 | 1.07 ± 0.26 | F(2,14)=1.281 p=0.3084 |
| *5-HT1B* | 1.02 ± 0.08 | 1.10 ± 0.17 | 0.85 ± 0.06 | F(2,15)=1.177 p=0.3352 |
| PFC | | | | |
| *5-HT1A* | 1.02 ± 0.08 | 1.15 ± 0.07 | 1.02 ± 0.08 | F(2,15)=0.8851 p=0.4332 |
| *5-HT1B* | 1.01 ± 0.07 | 1.06 ± 0.09 | 1.17 ± 0.17 | F(2,15)=0.4864 p=0.6242 |
| *5-HT2A* | 1.02 ± 0.07 | 1.01 ± 0.08 | 1.18 ± 0.16 | F(2,15)=0.7104 p=0.5072 |
| *5-HT4* | 1.00 ± 0.05 | 0.92 ± 0.07 | 0.86 ± 0.10 | F(2,14)=0.8139 p=0.4630 |
